# Comparative fMRI reveals differences in the functional organization of the visual cortex for animacy perception in dogs and humans

**DOI:** 10.1101/2024.11.12.623268

**Authors:** Eszter Borbála Farkas, Raúl Hernández-Pérez, Laura Veronica Cuaya, Eduardo Rojas-Hortelano, Márta Gácsi, Attila Andics

## Abstract

The animate-inanimate category distinction is one of the general organizing principles in the primate high-level visual cortex. Much less is known about the visual cortical representations of animacy in non-primate mammals with a different evolutionary trajectory of visual capacities. To compare the functional organization underlying animacy perception of a non-primate to a primate species, here we performed an fMRI study in dogs and humans, investigating how animacy structures neural responses in the visual cortex of the two species. Univariate analyses identified animate-sensitive bilateral occipital and temporal regions, non-overlapping with early visual areas, in both species. Multivariate tests confirmed the categorical representations of animate stimuli in these regions. Regions sensitive to different animate stimulus classes (dog, human, cat) overlapped less in dog than in human brains. Together, these findings reveal that the importance of animate-inanimate distinction is reflected in the organization of higher-level visual cortex, also beyond primates. But a key species difference, that neural representations for animate stimuli are less concentrated in dogs than in humans suggests that certain underlying organizing principles that support the visual perception of animacy in primates may not play a similarly important role in other mammals.

## Introduction

General principles that govern the organization of the high-level visual cortex as well as several functional specializations were described in visually oriented, highly social primate species (1–5). The animate-inanimate category distinction is one of the general organizing principles (1, 2, 6, 7), resulting in widely distributed animacy representations in the human high-level visual cortices. These representations show various diagnostic features of animacy, including mid-level features such as curvature, face-like and body-like features, and features reflecting capacity for movement and agency (8–10). The animate-inanimate distinction is also represented in homologous cortical areas of non-human primates (11).

Much less is known about the functional organization of non-primary visual areas of non-primate mammals, for which vision is secondary to other sensory modalities, with its significance not having increased as dramatically during evolution as for certain primates (3). For non-primates that are known to use vision for social interaction processing, behavioral evidence suggests that animacy cues are relevant. Cats and dogs distinguish biological motion from other forms of motions (3, 12, 13) and other animacy cues such as self-propulsion and speed changes can lead to an orienting response in dogs (14). When viewing natural scenes, dogs focus their gaze on animate entities, especially on their heads, more than on inanimate entities (15). Although recent neuroscientific works suggest a handful of high-level visual specializations in those non-primate species that primarily use vision for social interaction processing (3), data on how the representations of animate and inanimate entities are organized in non-primate visual cortices remains scarce.

Neuroimaging studies of non-primates directly comparing the visual processing of animate and inanimate entities have so far only been conducted with dogs. Dogs that live as companions constitute a special case for comparison to humans, even with their lower visual acuity and different color vision (16): dogs’ visual social environment is shared with humans, and they are the only non-human species that can be tested in an awake, unrestrained state (17). Three recent fMRI studies compared dog’ and humans’ high-level visual functions related to different aspects of animacy but remained inconclusive. Phillips and colleagues (18) found that a classifier distinguishing representations of dynamic animate and inanimate objects in humans failed to make the same distinction in dogs. In contrast, using static images, Boch and colleagues (19) identified body sensitivity as a factor accounting for animacy organization in the dog visual cortex. Finally, looking for representations of another diagnostic feature of animacy, faces, Bunford, Hernández-Pérez and colleagues (20) found areas exhibiting sensitivity for dog and human faces in humans but not in dogs. Together, these studies indicated that, similarly to the human brain, the dog brain exhibits sensitivity for certain aspects of animacy, but its extent and similarity to animacy-sensitivity in humans is still unclear.

To better understand the similarities and differences in how animacy structures higher-level visual perception in evolutionarily distant mammal species, we conducted a directly comparative dog-human fMRI study, in which participants viewed short videos of moving animate (dog, human, cat) and inanimate (car) entities. We selected stimuli to be therefore relevant to both species, and dynamic to involve motion-based features of animacy. To test animacy-sensitivity, we measured and compared neural response patterns elicited by these different stimulus classes. We predicted that if the animate-inanimate distinction is one of the general organizing principles of the dog visual cortex as it is in humans, we will find stronger responses to animate stimuli, and distinct response patterns to animate and inanimate stimuli, in non-primary visually responsive areas in the dog brain. Furthermore, we expected that if specific animate stimulus classes share some animacy features that are central for the functional organization of the human but not of the dog visual cortex – perhaps because dogs’ visual-social perception is far less detailed than humans’ –, then the neural responses for dog, human and cat stimuli would be less similar to each other and therefore overlap less in dogs than in humans.

## Methods

### Participants

15 family dogs (average (range) age of 7.1 (3-12.6); 7 spayed females, 1 intact-, 7 neutered males), and 13 humans (average age (range) of 31.5 (21-42), 7 females, 6 males) participated in the experiment. All dogs (7 Border Collies, 2 Golden Retrievers, 1 Australian Shepherd, 1 Labradoodle, and 4 mixed breeds) were mesocephalic and were trained to remain still inside the scanner (21). All human participants had normal or corrected to normal visual acuity and participated voluntarily. Dog owners and humans were recruited from the participant pool of the Department of Ethology at Eötvös Loránd University in Budapest, Hungary. All procedures were in accordance with relevant guidelines and regulations. Procedures involving dogs were approved by the Food Chain Safety and Animal Health Directorate Government Office, Hungary. Procedures involving humans were approved by the Committee of Scientific and Research Ethics (ETT-TUKEB), Hungary. Dog owners and human participants signed an informed consent.

### Experimental Design

The stimuli consisted of natural videos of dogs, humans, cats, and cars, some showing the whole body or object, some only parts, through close-up or wider shots (Figure 1 and Supplementary Video S1). The dog (D), human (H), and cat (C) videos constituted the animate (A) stimuli. We selected these animate stimulus classes to have an identical stimulus set for dogs and humans that involves familiar species to maximize the attention of dog subjects: the conspecific and two heterospecific species. Cars were selected as inanimate (iA) stimuli that move, vary in color and shape and all dog participants had ample experience with them. A reason to use a restricted number of stimulus classes was ensuring enough repetitions of each condition to elicit robust neural response in dogs where representations of categories are not established. All the videos were downloaded from YouTube (available under a Creative Commons License), cut to match the length required and resized to a final resolution of 1024 × 576 pixels. The videos displayed various situations and backgrounds using natural colors to minimize the possibility of systematic differences in visual properties across conditions and maximize ecological validity. To balance the differences in visual properties, brightness, contrast, hue, saturation, and motion for every video was calculated and videos were selected so that there were no significant differences in these properties between conditions in any of the runs (one-way ANOVA, ps > 0.05). To account for other low-level visual properties, captured by Gabor filters, the within- and between-category correlation of the outputs of the HMAX model’s C1 units (22) were compared and no differences were found for any of the categories (two-sample t test, ps > 0.05).

**Figure 1.**
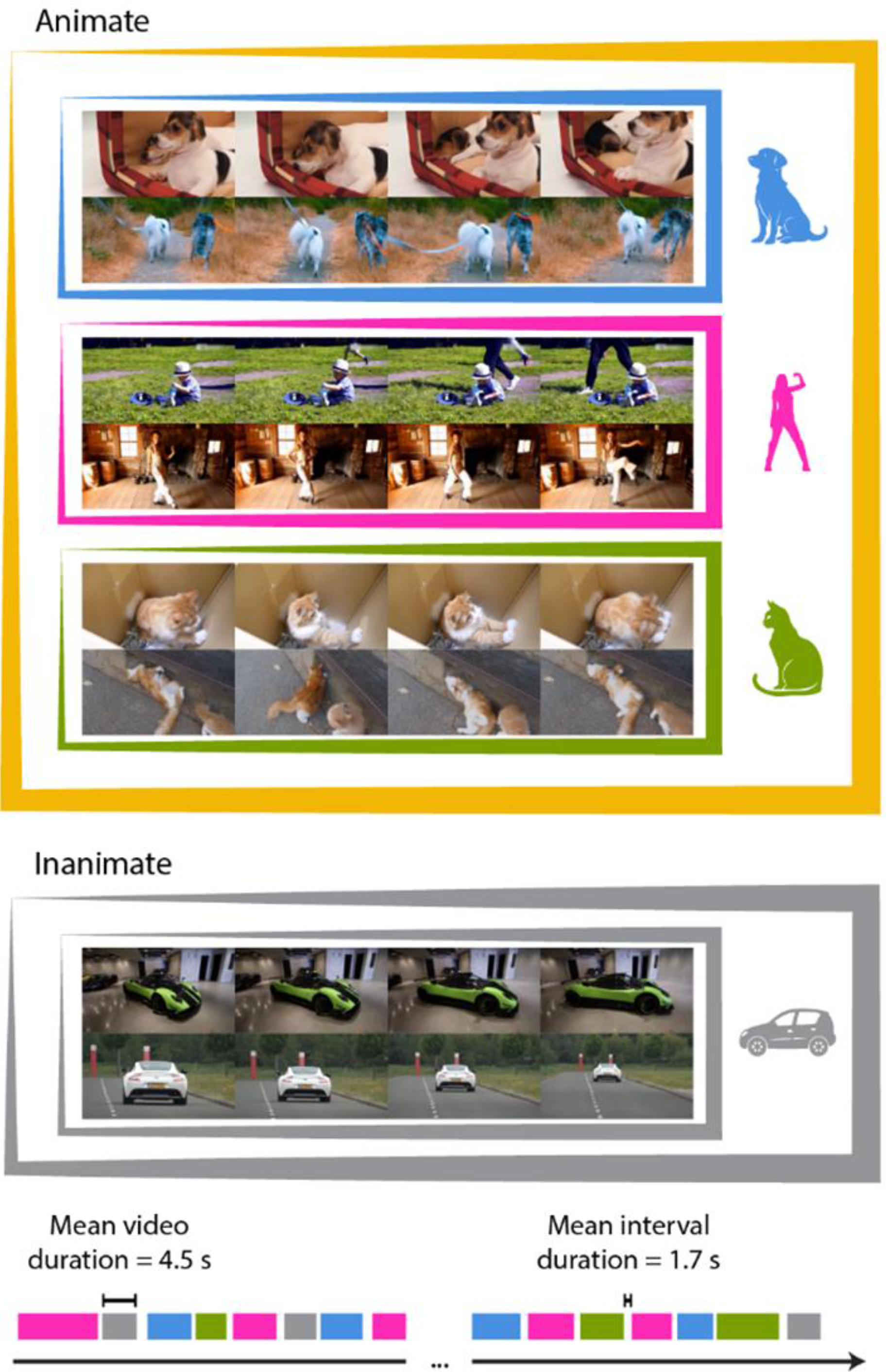
Experimental paradigm. Still images from two sample video stimuli representing each of the three animate (yellow) conditions: dog (blue), human (pink), cat (green) and the inanimate condition: car (grey) and the design of a single run. Each 310 s long run was composed of 12 videos (3.5-5.5 s, mean 4.5 s) of each of the four conditions followed by an interstimulus interval (0.7-2.5 s, mean 1.7 s). Stimulus design was identical for dog and human subjects. See also Video S1.

We used an event-related design (Figure 1). Each run contained different stimuli, 12 exemplars from each of the four conditions. The length of the videos varied between 3.5 and 5.5 s (mean 4.5 s). Each video was preceded by a grey screen with variable duration of 0.7 - 2.5 s (mean 1.7 s). The length and variability of stimulus duration and interstimulus interval were chosen to minimize predictability but maximize signal quality (23, 24). Runs ended with an additional 10 s of grey screen and lasted 310 s. The order of the videos within a run was randomized for every participant separately. Thirteen dogs completed six runs and two dogs completed only three, all humans completed six runs. Dogs were tested in two runs per session with rest periods of at least 20 min between sessions and a maximum of four runs per day. The dogs’ eyes were observed through an MR-compatible camera during the run to confirm they were awake during scanning. Four runs were discarded and repeated due to the dog falling asleep. Humans were tested in one or two days and in one session with 3 to 6 runs per day depending on the availability of the scanner and the participant.

Videos were presented using an MR compatible NordicNeuroLab LCD Monitor and controlled using Psychophysics Toolbox. Dogs viewed the screen directly, while humans viewed it through a mirror. The screen was set up at 155 cm from the head of the participants; the videos covered the entire height (49.8 cm) and width (88.6 cm) of the screen.

### fMRI Data Acquisition and Preprocessing

Data acquisition was performed at the Medical Imaging Centre at the Semmelweis University on a Philips 3T Ingenia scanner. For dogs, a Philips dStream Pediatric Torso 8ch coil and for humans, a Philips dStream Head 32ch coil was used. Functional data were collected using a gradient-echo-planar imaging (EPI) sequence (dogs and humans: TR=2500 ms, TE=20 ms, flip angle=90°, 2.5 mm-thick slices with 0.5 mm gap; dogs: field of view: 200×150×120 mm, acquisition matrix 80×58; 40 axial slices; humans: field of view: 240×240×122 mm, acquisition matrix 96×94; 41 axial slices). 124 volumes were acquired in each run. Anatomical data was collected using a T1-weighted 3D TFE sequence, with 1×1×1 mm resolution (dogs: at a separate session; humans: the end of the last functional imaging session). We acquired 180 slices covering the whole brain for anatomical localization.

Image preprocessing was conducted with FSL version 4.19 (Jenkinson et al., 2012). Statistical analysis of MRI data was performed using FEAT (FMRI Expert Analysis Tool) Version 6.00, part of FSL (FMRIB’s Software Library, www.fmrib.ox.ac.uk/fsl), time-series statistical analysis using FILM (25), and representational similarity analysis using PyMVPA software package (26) and MATLAB (2018b) custom code.

Volumes were motion-corrected and filtered using a 128 s high-pass filter. Scan-to-scan movement was calculated using framewise displacement (FD) (27), all the volumes where FD exceeded 1 mm were excluded (dogs 0.17%; humans 0%). The mean FD across subjects and runs was 0.07 mm for dogs and 0.05 mm for humans. Dog and human images were skull-stripped.

For dogs, we calculated a mean functional image that comprised all the volumes of all the runs for a given subject. This mean functional image was then manually transformed into a dog anatomic template (28) using 12 degree-of-freedom affine transformation, and the transformation matrix was used to transform all the volumes into the template space. Human images were first registered to the anatomical scan using a 7 degree-of-freedom linear transformation, then to the Montreal Neurologic Institute MNI template using 12 degree-of-freedom affine transformation. Next, images were smoothed (Gaussian kernel, FWHM 5 mm) for both dogs and humans.

### fMRI Statistical Analysis

#### RSA Comparing Neural and an EVC Model

To map neural responses to low-level visual properties, we assessed, using RSA, where the representational geometry of the stimuli in the dog and human brains was similar to how an early visual cortex (EVC) model represent them.

First, representational dissimilarity matrices (DSM) of the stimulus set were calculated to characterize the pairwise dissimilarity of the response patterns elicited by the stimuli (Kriegeskorte, 2009). For this, each stimulus event was modelled convolving its times series with the canonical hemodynamic response function and the resulting, normalized β values were used to describe every voxel by how dissimilar a sphere of voxels (r=2 voxels for dogs and r=3 voxels for humans) around it represents each pair of the stimuli. Dissimilarity was measured as the correlation distance (1 - Pearson correlation). DSMs were calculated this way for each run of every participant and then were averaged across runs and participants. In the case of dogs, this process was repeated for the whole brain. In the case of humans, we restricted the analysis to the occipital and temporal lobes (delineated based on MNI atlas) where the visual areas are, to reduce the computation load.

Second, the stimuli were presented to the computational model HMAX (22). In its first and second layers, the HMAX model simulates the selective tuning of neurons in primary visual cortex (V1) and has been shown to match with the activity patterns of the EVC of the primate brain (11, 29, 30). Specifically, we used the second layer of the model (referred here as EVC model), which corresponds to V1 complex cells. This layer yields a response vector of a V1 unit by performing a 2-dimensional convolution of a Gabor filter with an image. The algorithm was applied to each frame of each video using a set of Gabor filters with four different orientations (−45°, 0°, 45°, 90°) and 17 sizes (7 to 39 pixels in 2-pixel steps) per orientation, modeling 68 V1 units for each frame of a video. To obtain a unidimensional response vector for every video stimulus, principal component analyses were performed within the response vectors of every V1 unit across all frames and for all stimuli. The first principal components were used in the following step, because those contained the most explained variance for the given video. Next, the DSM of response vectors was calculated to assess the representational geometry expected based on our stimuli’s low-level visual features.

Third, to compare the brain representation and the model, correlation maps were created, in which each voxel was assigned the Pearson correlation between neural DSM and model DSM. To account for the spatial dependency of the searchlight approach (31) and to reduce the number of assumptions on the maps, we used a random permutation method (32) for group-level analysis and statistical inference.

The stimulus labels were randomly swapped, a random DSM map and then the correlation map between the random DSM map and the DSM of the model were created. This process was repeated 1,000,000 times. Next, the maps were averaged to calculate the correlations expected by chance and their standard deviations. These parameters were then used to transform the original correlation maps to Z-score maps. The maps were then filtered to maintain only the voxels with a Z-score > 3.1 (equivalent to p < 0.001). The random correlation maps were also filtered at Z-score > 3.1. Using these maps, the cluster size distribution expected by chance was calculated and this distribution was used to filter out the clusters with a size smaller than the size expected by chance (p < 0.05).

#### General Linear Model (GLM)

To identify regions that are sensitive to our conditions, univariate analysis with the four conditions (D, H, C, iA) as regressors were conducted using the FSL’s implementation of the General Linear Model.

The first-level individual models included the movement parameters as regressors of no interest. The regressors were convolved with the canonical hemodynamic response function, then using all six runs, condition effects were estimated for each participant. At the group level, a random-effects analysis was performed for each species.

For further analyses, the visually responsive regions of the dog and human cortex were identified by comparing all conditions to the implicit baseline. To determine the brain regions sensitive to animate stimuli, the videos from all three animate conditions (D, H, C) were grouped and contrasted with the inanimate condition (A>iA contrast). To further characterize the animacy-sensitivity of the dog and human brains, each of the three animate conditions was contrasted with the inanimate condition separately (D>iA, H>iA, C>iA, specific contrasts).

For all contrasts, the resulting Z-statistic (Gaussianised T/F) images were thresholded at Z > 3.1 and the threshold for cluster-correction was p = 0.05 (Worsley, 2001).

##### Response profiles

To further characterize the A>iA contrast results, for every subject and for each group level-derived peak, first the parameter estimates (β weights) for each condition in a sphere (radius = 3 voxels) centered around the peak was assessed using FSL’s Featquery tool. Second, these parameter estimates were compared on the group level, using t tests.

##### Overlap calculation

To determine and compare across species the extent to which specific contrast maps overlap, first, the visually response cortex was determined in both species by contrasting all conditions to baseline using a cluster threshold of Z>3.1 and a corrected cluster significance threshold of p=0.05. Then, the number of suprathreshold voxels were assessed for each specific contrast as well as the number of voxels in the visually responsive cortex in dogs and in humans. Then, the percentage of those visually responsive voxels that were suprathreshold in (1) all three specific contrasts (3-overlap) or (2) two of the three specific contrasts (2-overlap), were calculated, and the percentages of both 3-overlapping and 2-overlapping voxels were compared between dogs and humans, using t tests. *Species-preference*. We also assessed the proportions of animate-sensitive voxels that responded strongest to conspecific stimuli or to any of the heterospecific stimulus classes. For that, each animate condition (D, H, C) was contrasted with the implicit baseline, and the average Z-scores per contrast were calculated across all dog and across all human participants in every animate-sensitive voxel. The condition from the contrast with the highest average Z-score was assigned as the preferred species for each voxel. To determine whether the percentage of voxels differed above chance (0.33) for the three conditions, permutation test was performed by randomly swapping the labels of the stimuli, calculating the preference for each voxel and their percentages 1,000,000 times.

#### Category- and Class-boundary Effect Tests

To identify regions with greater representational dissimilarity for between-category (i.e., animate-inanimate) than within-category (i.e., animate-animate) stimulus pairs (category boundary effect test as described in Kriegeskorte et al., 2008), we calculated DSMs per run, per participant for both between-category and within-category pairs across the brain (for dogs: whole brain, for humans: temporal and occipital cortices), using a searchlight approach (sphere with r=2 voxels for dogs, and r=3 voxels for humans). DSMs were then averaged across runs and participants, and a difference map was calculated. To determine whether the difference was significant at the voxel level, we repeated the process 1,000,000 times but randomly swapping stimulus labels, thus generating 1,000,000 permuted difference maps. We transformed each voxel of the original difference map to Z-score by comparing it with the corresponding voxel in the permuted difference maps, generating a Z-score map. To determine cluster size threshold, we repeated the same process for the permuted difference maps, thus generating 1,000,000 permuted Z-score maps. All Z-score maps were thresholded at Z > 3.1. We calculated the cluster size distribution in the permuted Z-score maps and used it to determine the cluster size expected by chance in the original Z-score map. All clusters with a size smaller than expected by chance (p<0.05) were excluded. Category-boundary tests were performed in a similar way for between-class (e.g., dog-car) vs. within-class (e.g., dog-dog) stimulus pairs.

##### Overlap calculation

To determine and compare across species the extent to which class-boundary tests overlap, first, the results were binarized. Then, the percentage of the voxels active in (1) all three classes (3-overlap) or (2) two of the three classes (2-overlap), were calculated, and the percentages of both 3-overlapping and 2-overlapping voxels were compared between dogs and humans, using t tests.

## Results

### RSA Comparing Neural and an EVC Model

For results in dogs and humans, see Figure 2 and Supplementary Table S1. We found a significant correlation with the early visual cortex model in clusters in the occipital cortex of dogs and humans. In dogs, the cluster in the right hemisphere was centered in the marginal gyrus (MG) and the cluster in the left hemisphere was centered in the splenial gyrus (SpG). In humans, the cluster extended bilaterally, with main peaks in the bilateral calcarine fissure (CAL).

**Figure 2.**
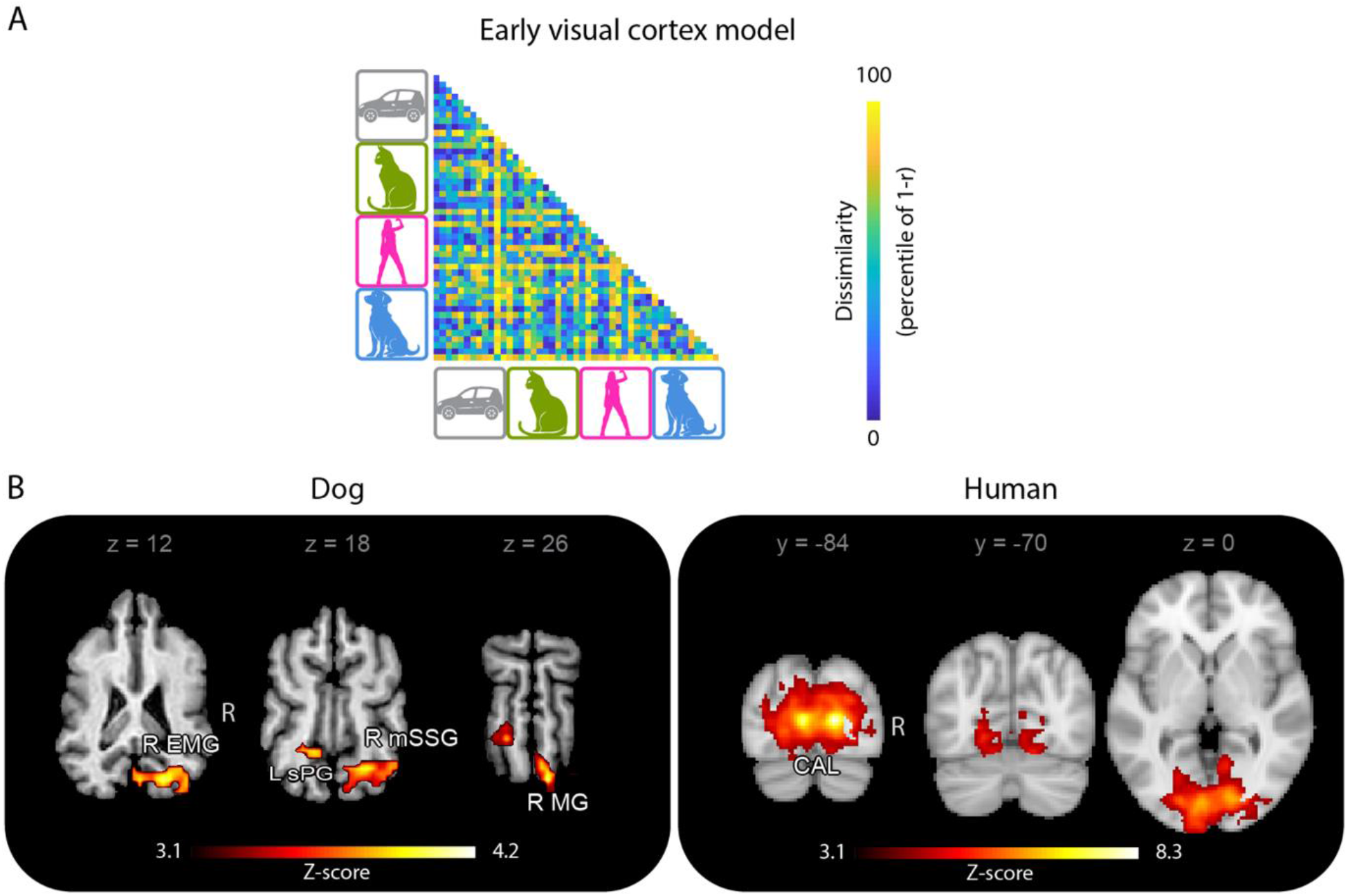
RSA comparing neural and EVC model representations. A, DSM for the EVC model. B, Results for correlation with the EVC model in dogs (n = 15) and humans (n = 13) superimposed on a template brain (for dogs: Czeibert et al., 2019; for humans: MNI template brain), using a cluster threshold of Z>3.1 and a corrected cluster significance threshold of p=0.05 in a searchlight analysis (sphere radius = 3 voxels for dogs and 3 voxels for humans). L=left; R=right; EMG=ectomarginal gyrus; SpG=splenial gyrus; mSSG=mid suprasylvian gyrus; MG=marginal gyrus; CAL=calcarine fissure and surrounding cortex.

### GLM

For visual responsiveness (all stimuli > implicit baseline) results in dogs and humans, see Supplementary Table S2. For GLM results for each contrast (A>iA, D>iA, H>iA, C>iA) in dogs and humans, see Figure 3 and Supplementary Table S3. In both species, these contrasts revealed clusters that extended mainly through the temporal and occipital lobes. Specifically, in dogs, for A>iA, we found bilateral clusters in the mid suprasylvian gyrus (mSSG), extending caudally in the left hemisphere, and bilateral clusters in the ectomarginal gyrus (EMG); for D>iA, left SpG extending to mSSG, and a right cluster in the suprasylvian gyrus (SSG); including mid and caudal portions; for H>iA, left clusters including the mid ectosylvian gyrus (mESG), the mSSG, and the SpG, and a right cluster including mid and caudal portions of the SSG; and for C>iA, a left cluster in the caudal part of the SSG. In humans, for all four contrasts, we found clusters bilaterally, involving portions of the inferior temporal gyrus (ITG), middle temporal gyrus (MTG), inferior occipital gyrus (IOG) and the fusiform gyrus (FFG), typically extending more to the temporal lobe in the right hemisphere. Activity response profiles for GLM-derived A>iA peaks in dogs and humans are shown in Figure 3B.

**Figure 3.**
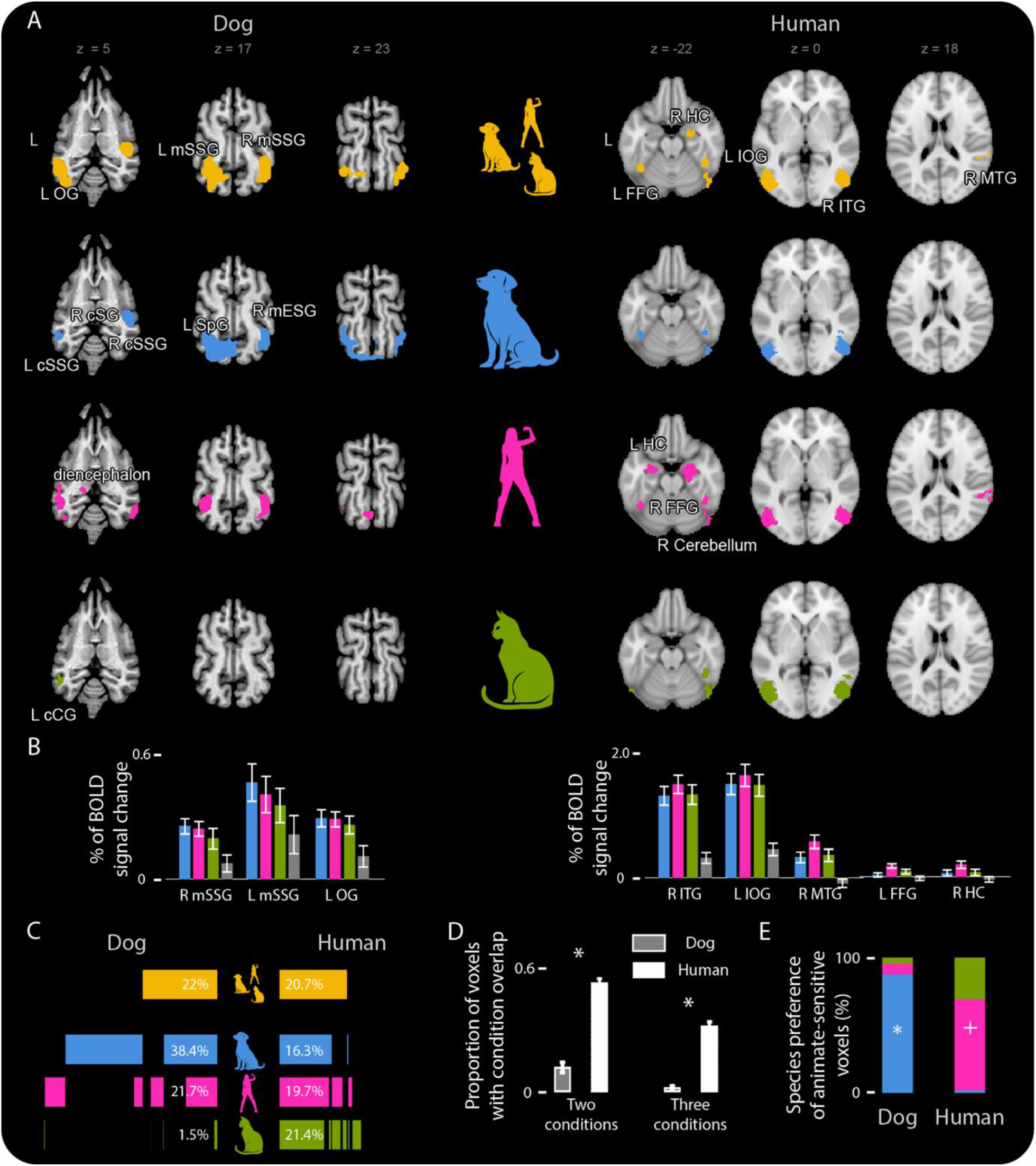
GLM results in dogs and humans. A, Dog and human contrast maps superimposed on a template brain, using a cluster threshold of Z>3.1 and a corrected cluster significance threshold of p=0.05. Upper row: A > iA (yellow), lower rows: D > iA (blue), H > iA (pink), C > iA (green). B, Response profiles. Bar graphs represent parameter estimates (beta weights) in select GLM-derived peaks (sphere radius = 3 voxels) for each condition. C, Activity proportions and overlaps. Horizontal bars (and percentages) represent the proportions of suprathreshold voxels per condition, relative to the total voxel number, within the visually responsive cortex; vertical overlaps represent activity overlaps. D, Overlap calculation results. The percentage of suprathreshold voxels within the visually responsive cortex for which at least two conditions overlap and in which all three overlap. E, Species-preference. Proportion of animate-preferring voxels for which the strongest response was found for the given condition. L=left; R=right; OG = occipital gyrus; mSSG = mid suprasylvian gyrus; cSSG = caudal suprasylvian gyrus; cSG = sylvian gyrus; SpG = splenial gyrus; mESG = mid ectosylvian gyrus; cCG = caudal composite gyrus; FFG = fusiform gyrus; HC=hippocampus; IOG = inferior occipital gyrus; ITG=inferior temporal gyrus; MTG = middle temporal gyrus. *: p<0.01, +: p=0.062. Error bars represent SE.

#### Overlap calculation

Comparing the proportion of overlapping activities within the visually responsive cortex, we found that in the dog brain there was a significantly lower proportion of voxels than in the human brain in which all three (D, H, C) contrast maps overlapped at z=3.1 (dogs: 2%, humans: 31%) (t(14.781) = 12.044, p < 0.01). The proportion of overlapping voxels between at least two categories out of the three was also lower in the dog brain compared to that of the human brain (dogs: 11%, humans: 51%) (t(25.925) = 11.236, p < 0.01) (Figure 3D).

*Species-preference*. Analyses of the extent to which voxels in the animate-inanimate contrast responded stronger to dog or human or cat stimuli indicated that in dogs, the majority of animate-preferring voxels (87%) responded strongest to conspecific (dog) stimuli (likelihood of obtaining the observed proportions by chance, using permutation testing: p = 0.004). In humans, the majority of animate-preferring voxels (67%) responded strongest to conspecific (human) stimuli (permutation testing confirmed a marginally significant effect, p = 0.062) (Figure 3E).

### Category- and Class-boundary Effect Test

For category- and class-level boundary test results in dogs and humans, see Figure 4 and Supplementary Table S4. We found temporal and occipital clusters, located similarly to those identified by univariate animacy-sensitivity contrasts. Specifically, in dogs, for A>iA, we found bilateral mSSG/mESG clusters; for D>iA, a left cluster centered in the caudal suprasylvian gyrus (cSSG) and an mSSG-centered right cluster; for H>iA, left clusters in cSSG and caudal ectosylvian gyrus (cESG), and right clusters in EMG and rostral sylvian gyrus (rSG); and for C>iA, a left cluster in mESG, and a right SpG cluster, both extending caudally. In humans, for all contrasts, we found large clusters in both hemispheres, extending from MTG to ITG and IOG, and ventrally to FFG, with similar peaks across contrasts.

**Figure 4.**
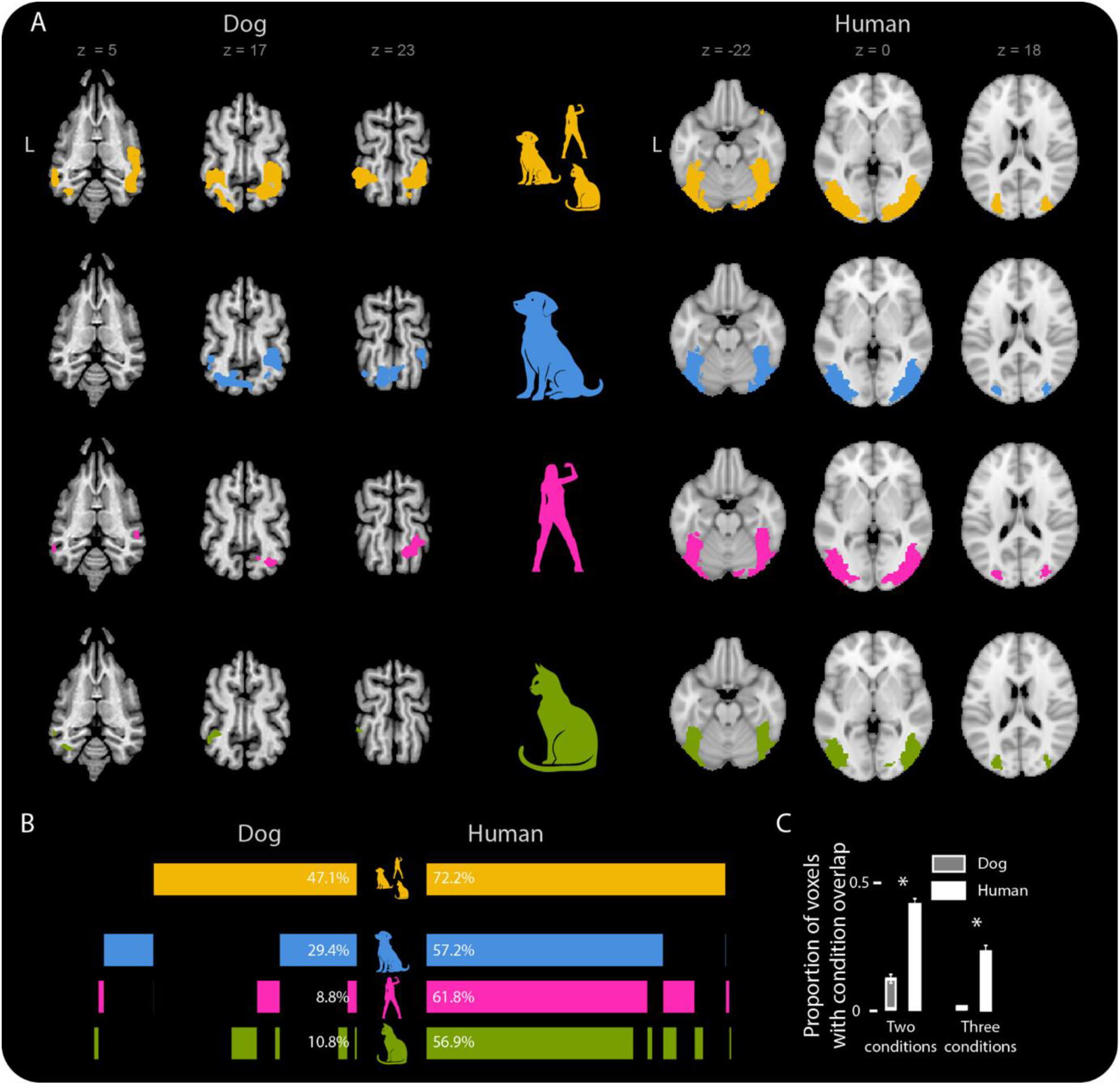
Category boundary test results in dogs and humans. A, Dog and human results superimposed on a template brain using a cluster threshold of Z>3.1 and a corrected cluster significance threshold of p=0.05. B, Activity proportions and overlaps. Horizontal bars (and percentages) represent the proportions of suprathreshold voxels per condition, relative to the total voxel number, within the visually responsive cortex; vertical overlaps represent activity overlaps. C, Overlap calculation results. The percentage of suprathreshold voxels within the visually responsive cortex for which at least two conditions overlap and in which all three overlap. L=left. *: p<0.01. Error bars represent SE.

#### Overlap calculation

Proportion of overlapping voxels within the visually responsive cortex were significantly lower in dogs than in humans, both in case all three (D, H, C) similarity maps overlapped (dogs: 1%, humans: 26%) and in case at least two of the three similarity maps overlapped (dogs: 14%, humans: 47%), (t(12.217) = 8.518, p < 0.01 and t(20.981) = 9.405, p < 0.01) (Figure 4C).

## Discussion

This study investigated how animacy structures neural responses in the visually responsive cortex of dogs and humans. We found both similarities and differences across the two species. Univariate analyses identified bilateral occipital and temporal regions in both species responding stronger to animate than inanimate stimuli. These animate-sensitive regions are distinct from functionally determined early visual areas (i.e., that represented the stimuli similarly to an EVC model). Most animate-sensitive peaks preferred all animate stimulus classes over inanimate stimuli, but more animate-sensitive voxels responded stronger to conspecific than to any of the heterospecific stimuli in dogs, revealing conspecific-preference. A similar trend was observed in humans. The areas sensitive to the specific class of animate stimuli, dog, human and cat, overlapped less in dog than in human brains. Multivariate tests revealed regions in both species in which the representational geometry of stimulus pairs within animate and within inanimate categories were more similar than that of stimulus pairs crossing the animate-inanimate boundary. Such boundary effects were identified in both species for dog, human and cat stimuli as well, and these overlapped less in dogs than in humans. The regions exhibiting these categorical representations for animate stimulus classes largely overlapped with univariate animacy-sensitive clusters.

The similarities between dog and human neural response patterns, namely animacy-sensitivity and category boundary effects in visual cortical regions, demonstrate that animacy is an organizing principle in the visual perception of both species, and the representations of animate entities exhibit categorical structure. Categorical representation has already been shown across visual domains in multiple primate species. Going beyond these previous results, the present work suggests that animacy and the subordinate categorical feature of the input may determine the location and the structure of the neural response not only in primates’, but at least in some non-primate mammals’ visual perception too. We also note, however, that as animacy is multidimensional in nature (8–10), dogs’ and humans’ animacy sensitivity, as reported here, do not necessarily originate from the representation of the same animacy dimensions. Nevertheless, the present findings provide evidence that dogs’ visual cortex contains functionally organized areas, as primates’ does.

The animacy-sensitive regions reported here correspond to but extend beyond those reported previously (8, 9, 19). Specifically, for dogs, we found that animacy sensitivity characterizes 22% of the visually responsive cortex, including regions bilaterally in the EMG and mSSG. Using static stimuli, Boch et al. (2023) found similarly located, although less extensive regions as sensitive for animate (specifically, body and face) stimuli. Various factors may have contributed to this difference between studies, including the use of dynamic stimuli in the present study, as these stimuli also carried motion-related animacy cues (33, 34), but also scanning-technical factors. Prior results on dog neural responses to (human) faces vs. objects, potentially also driven by animacy sensitivity, also suggested the involvement of bilateral temporal regions (35, 36). For humans, animacy sensitivity characterized 20,7% of the visually responsive cortex and included extended regions in the higher-level visual cortex. Whereas previous works often focused on specific aspects of animacy (such as face, body, humanness, agency, biological motion), our highly natural stimuli carried multiple aspects of animacy at the same time, and this could have resulted in the finding of more extensive regions here (9, 11).

Beyond similarities, differences in animacy representations were also found between dogs and humans. The main species difference, that the detected neural activity evoked by animate stimulus classes overlapped less in dogs than in humans, may reflect how evolution drove mammalian brain organization for visual perception on a global level. Adaptive visual-social behavior and underlying neural representations shaped by relevant behavioral goals differ substantially between primates and (even visually oriented) non-primate mammals. The greater emphasis on visual social communication in primates, their more nuanced social interactions and within-face movements necessitate the efficient processing of bodily detail-encoding social cues and signals beyond body postures more than in non-primates. Indeed, in non-primates the relative role of perception and interpretation of whole-body actions is arguably greater. Therefore, the behavioral relevance of our stimuli showing moving dogs, humans and cats may have differed for human and dog subjects. Dogs’ focus during the visual perception of animate stimuli may be on bodily actions while humans’ rather on faces and gestures. This notion is supported by previous canine neuroimaging work: On one hand, these studies found no convincing evidence for the existence of face-sensitive areas in dogs, and body sensitivity, weak and restricted to small regions, has been reported for this species in a single study so far. On the other hand, Phillips et al. (18) found that representations of different actions of animate entities were separable in the dog brain, but the representations of animate and inanimate objects were not. Therefore, the pattern observed in the present study suggests that animacy sensitivity in the dog brain does not primarily stem from the common presence of face and body across all animate stimulus classes, but rather from the representation of biological motion. And it is possible that the representations of different biological motions in dogs are less generalized across the different animate stimulus classes than face and body representations in humans. These findings are in line with the proposal that dogs’ high-level visual cortex may contain distinct agent-responsive areas, and also with the specific sensitivities that have been described in these areas (Boch et al., 2024). Future research should assess the validity of this potential explanation behind the species differences in animacy perception.

The finding on conspecificity-preference in dogs, that the majority of animate-sensitive voxels responded maximally to conspecific stimuli, and a similar tendency in humans, corroborate and extend previous results. Recently, Bunford, Hernández-Pérez et al. (2020) identified conspecific-preferring clusters in the visually responsive cortex in both species. In that paper, in humans, conspecific-preference was restricted to face-sensitive regions. The present findings, using more naturalistic and more variable videos, support these previous results and suggest that conspecific-preference may be characteristic for animate-sensitive visual regions, at least in dogs. Conspecific-preference, therefore, may be present at different stages of visual processing of faces and animacy. This suggests that the bias for processing conspecific stimuli does not reflect a functional specialization but rather a more generalized preference, perhaps driven by attention or motivational relevance (37, 38).

This work has a few potential limitations. First, inanimate stimuli consisted of cars only, and animate stimuli included only a narrow but highly relevant subset of all animate entities that dogs and humans may encounter, which selections, for reasons detailed in the Methods may also have affected the generalizability of the animacy sensitivity results. Potential differences in the level of relevance between animate and inanimate conditions may have also been present: for example, human stimuli may have been more relevant than cars to dogs. The similarity of the cortical locations of our animacy sensitivity results to previous findings in both dogs (19) and humans (8, 9), however, suggests that it was indeed the presence of animacy in dog, human and cat stimuli (and its lack in car stimuli) that drove the measured differences in neural activity. Furthermore, although our stimuli did not involve a wide variety of animate classes, they, being videos, carried not only static but also action-related diagnostic features of animacy, in a highly natural manner.

Another potential limitation of any directly comparative visual study of dogs and humans may stem from their different visual capacities: dogs perceive less details of visual stimuli presented on a monitor than humans, due to their different color vision, lower critical flicker-fusion threshold, lower depth perception, and lower visual acuity [missing references]. As a consequence, dogs may have seen our stimuli as more similar than humans have, and this may have affected neural response patterns. While we cannot exclude such effects, we do not think that visual capacity differences between dogs and humans confounded the key species difference presented here, that is the smaller overlap of condition-specific clusters for dogs. On the one hand, it can be assumed that visual capacities influenced the perception of animate and inanimate stimuli similarly. On the other hand, the less detailed visual perception of dogs may lead to less differentiated and thus more overlapping representations of the conditions, relative to those in humans, but not to less overlapping ones, as was the case here.

In conclusion, the findings presented here suggest that the importance of the animate-inanimate distinction may be reflected in the organization not only of primate, but more generally of mammalian higher-level visual cortex. The key species difference, that neural representations for animate stimuli cluster less in dogs than in humans demonstrates that different animate stimulus classes may not form a unified category in dogs and suggests that the dog brain may lack those functional specializations that drive the unified response in humans. To understand the principles that organize the visual perception of animate objects in non-primate mammals, other factors need to be considered than those underlying the animacy organization of the primate visual cortex.

## Acknowledgements

This project was funded by the Hungarian Academy of Sciences [MTA Lendület (Momentum) Programme (LP2017-13/2017), National Brain Programme 3.0 (NAP2022-I-3/2022]; the Eötvös Loránd University; the European Research Council [European Union’s Horizon 2020 research and innovation program (grant number 950159)], the HUN-REN – ELTE Comparative Ethology Research Group (01 031). R.H.-P. and L.V.C. were supported by the Mexican Council of Science and Technology (CONACYT, 407590 and 409258, respectively). R.H.-P. was also supported by the Austrian Science Fund (FWF) (10.55776/ESP602).

We thank all dog and human participants and dogs’ owners for their participation and contribution to these data.

## Author Contributions

R.H.-P., L.V.C., E.B.F. and A.A. performed conceptualization. R.H.-P., L.V.C. performed investigation. R.H.-P. and E.R.-H. performed formal analysis. R.H.-P., E.B.F and A.A. performed visualization. E.B.F., R.H.-P. and A.A. performed writing – original draft. E.B.F., R.H.-P., M.G. and A.A. performed writing – review & editing.

## Supplementary information

**Supplementary Table S1.**

| Brain region | Cluster size (voxel) | Peak Z-score | Coordinates (x, y, z) |
| --- | --- | --- | --- |
| Dogs |  |  |  |
| R MG | 254 | 4,234 | 5; -35; 26 |
| R EMG |  | 3,675 | 11; -33; 10 |
| R mSSG |  | 3,462 | 21; -29; 22 |
| L SpG | 101 | 4,248 | -9; -25; 22 |
| Human |  |  |  |
| R CAL | 8006 | 6,414 | 14; -84; 6 |
| L CAL |  | 5,958 | -6; -86; 6 |
| R CUN |  | 4,343 | 16; -88; 22 |
| R LING |  | 4,196 | 14; -78; -10 |
| L LING |  | 4,189 | -20; -84; -2 |
| S SOG |  | 4,083 | -14; -82; 20 |
| L CAL |  | 3,937 | -8; -100; -2 |
| R MOG |  | 3,892 | 30; -88; 14 |
| L SOG |  | 3,841 | -20; -100; 16 |
| L CER6 |  | 3,800 | -18; -76; -18 |
| L LING |  | 3,788 | -22; -66; -4 |
| R LING |  | 3,714 | 12; -62; 0 |
| L CUN |  | 3,687 | 4; -96; 14 |
| R FFG |  | 3,646 | 30; -82; -8 |
| L MOG |  | 3,511 | -32; -84; 14 |
| L CAL |  | 3,494 | 6; -96; -12 |
| L LING |  | 3,430 | -2; -78; -8 |
| L CUN |  | 3,419 | 2; -88; 30 |
| R LING |  | 3,269 | 22; -98; -10 |
| R SOG |  | 3,243 | 26; -78; 30 |
| L MOG |  | 3,214 | -28; -98; 2 |
| L CAL |  | 3,209 | -18; -68; 12 |
| L CUN |  | 3,206 | -12; -82; 36 |

RSA results for dogs and humans. Threshold for reporting: cluster threshold of $Z > 3.1$ and a corrected cluster significance threshold of $p = 0.05$ . All peaks $\geq 16$ mm apart are reported. L=left; R=right; MG=marginal gyrus; EMG=ectomarginal gyrus; mSSG=mid suprasylvian gyrus; SpG=splenial gyrus; CAL=calcarine fissure and surrounding cortex; CUN=cuneus; LING=lingual gyrus; SOG=superior occipital gyrus; MOG=middle occipital gyrus; CER6=lobule VI of cerebellar hemisphere; FFG=fusiform gyrus.

**Supplementary Table S2.**
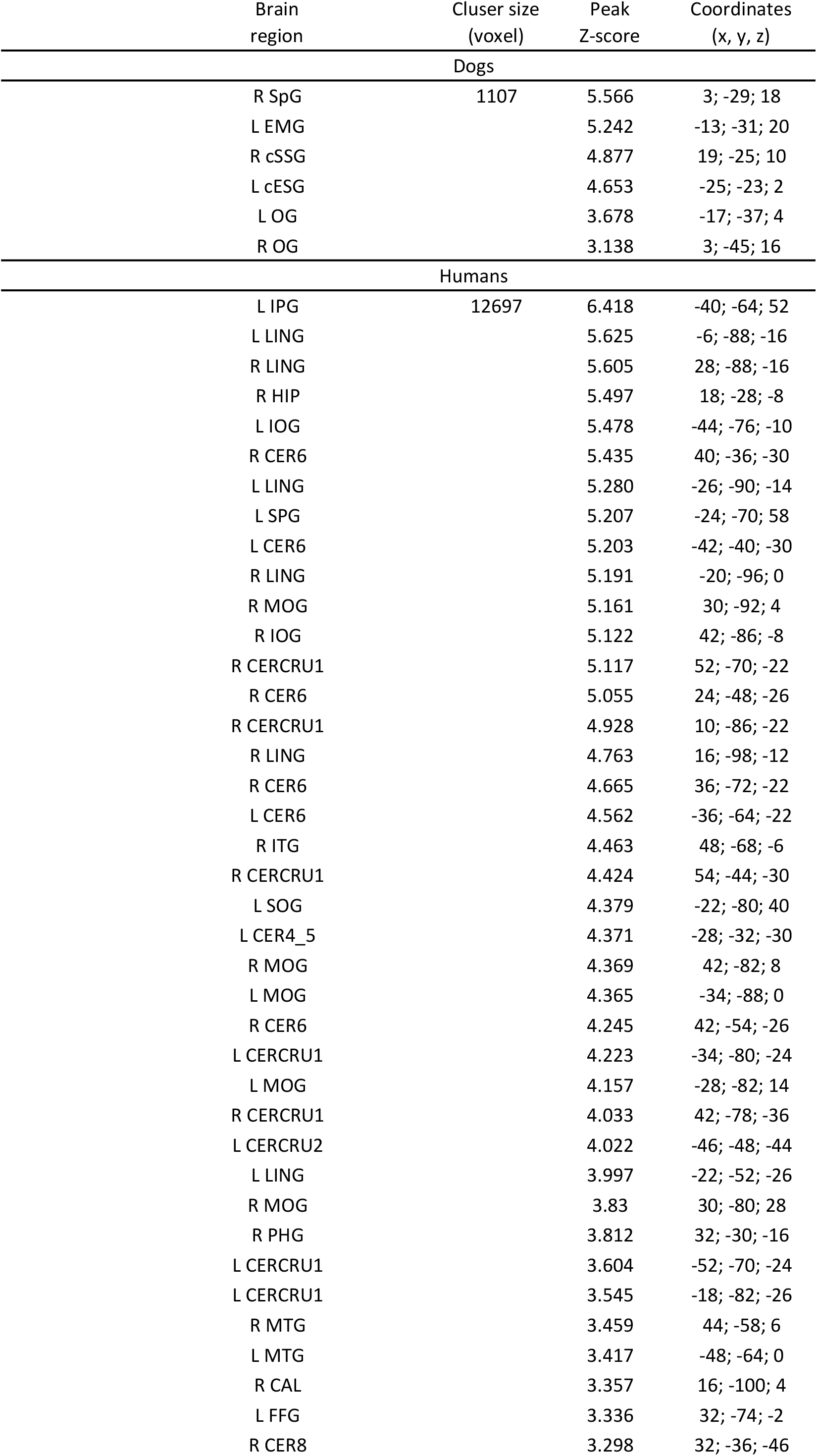
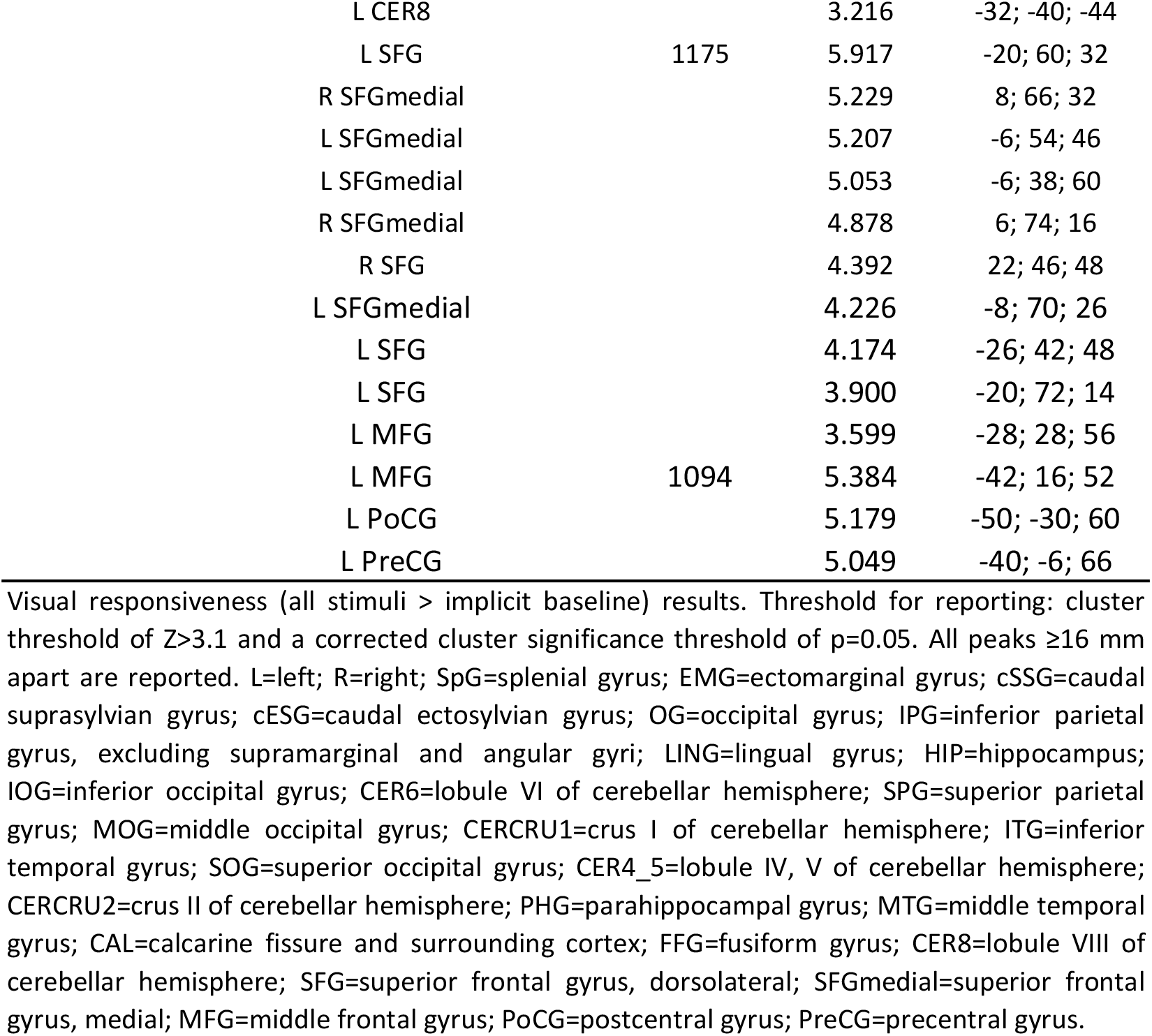

**Supplementary Table S3.**
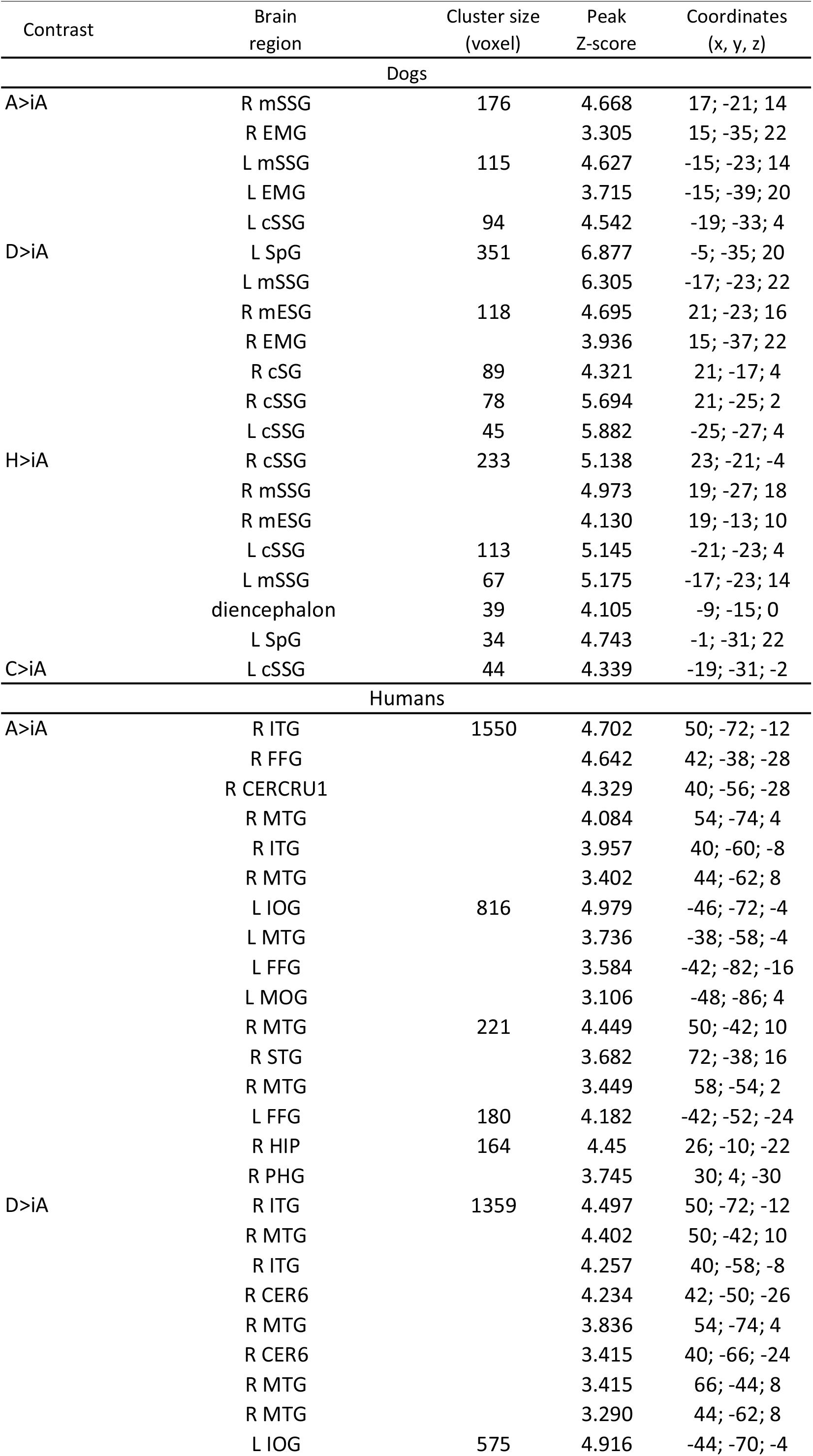
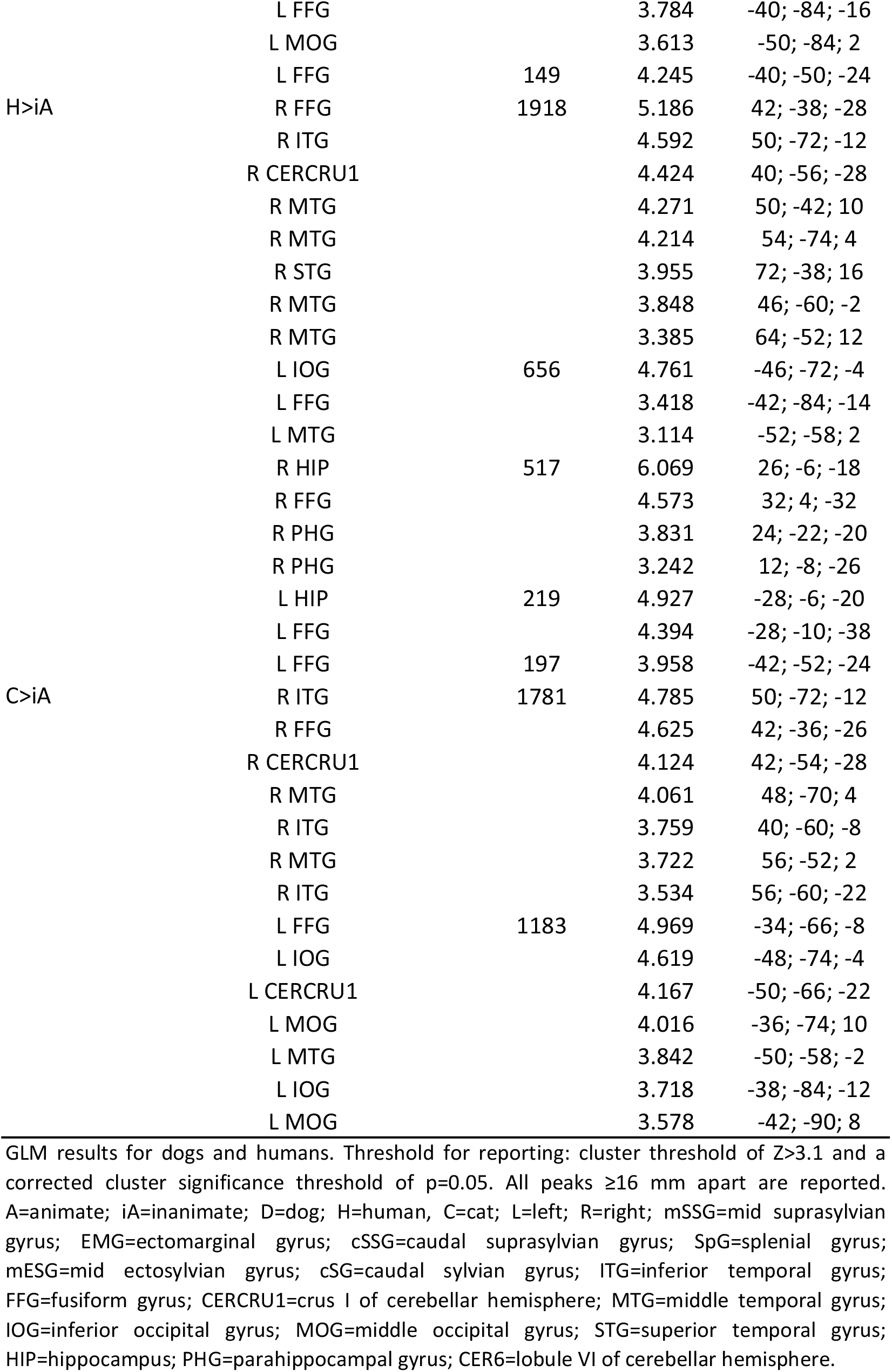

**Supplementary Table S4.**
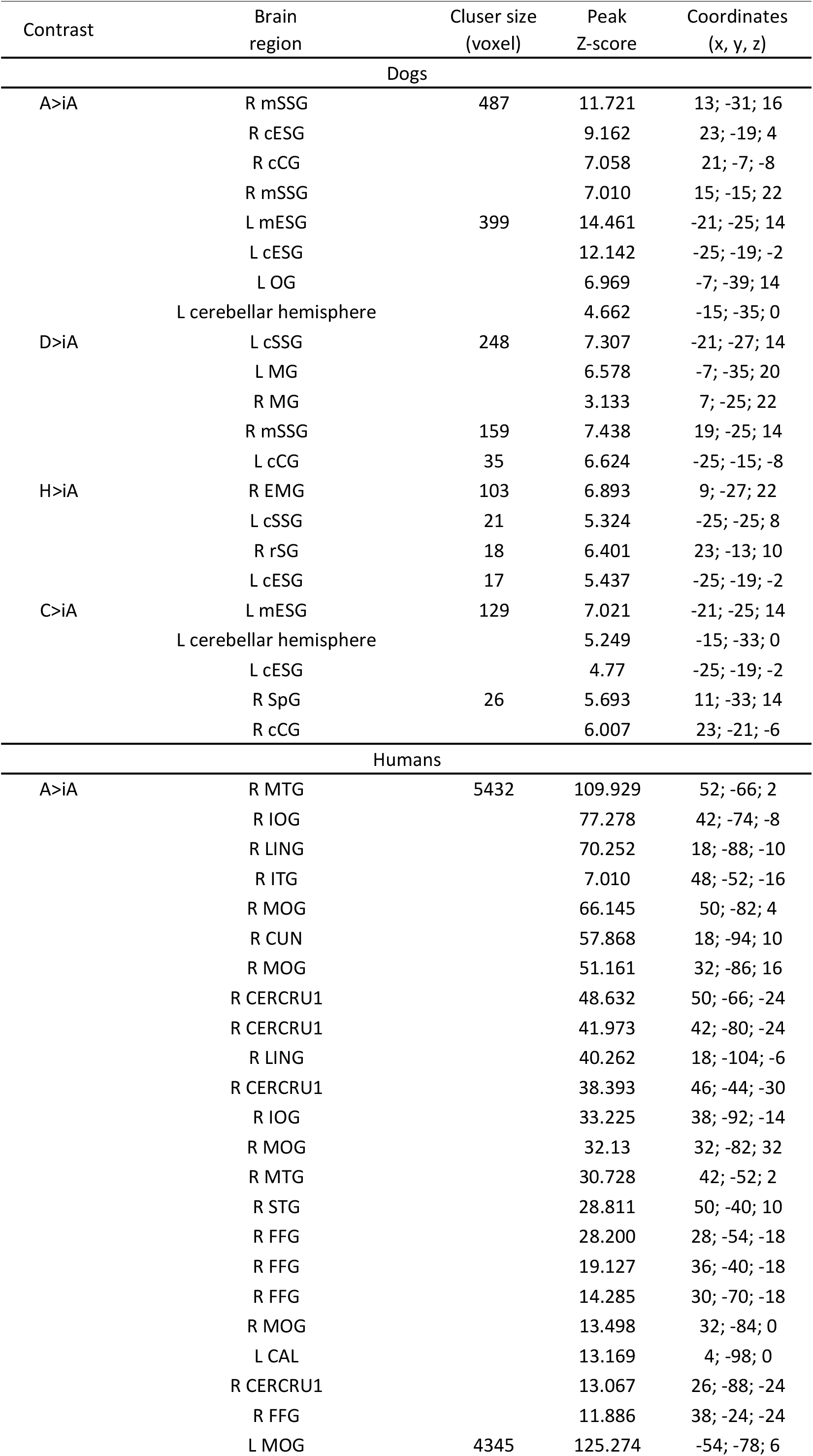
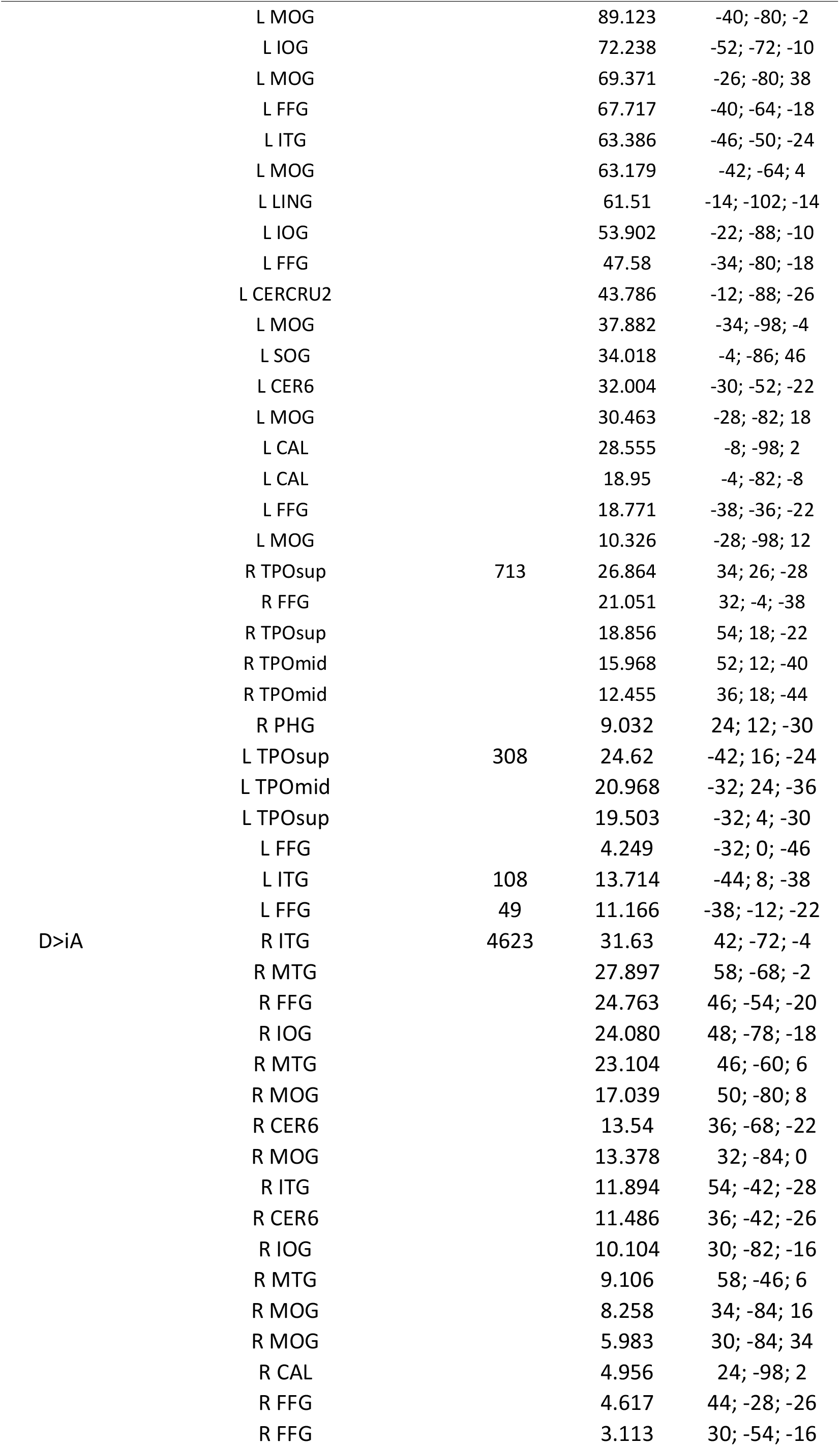
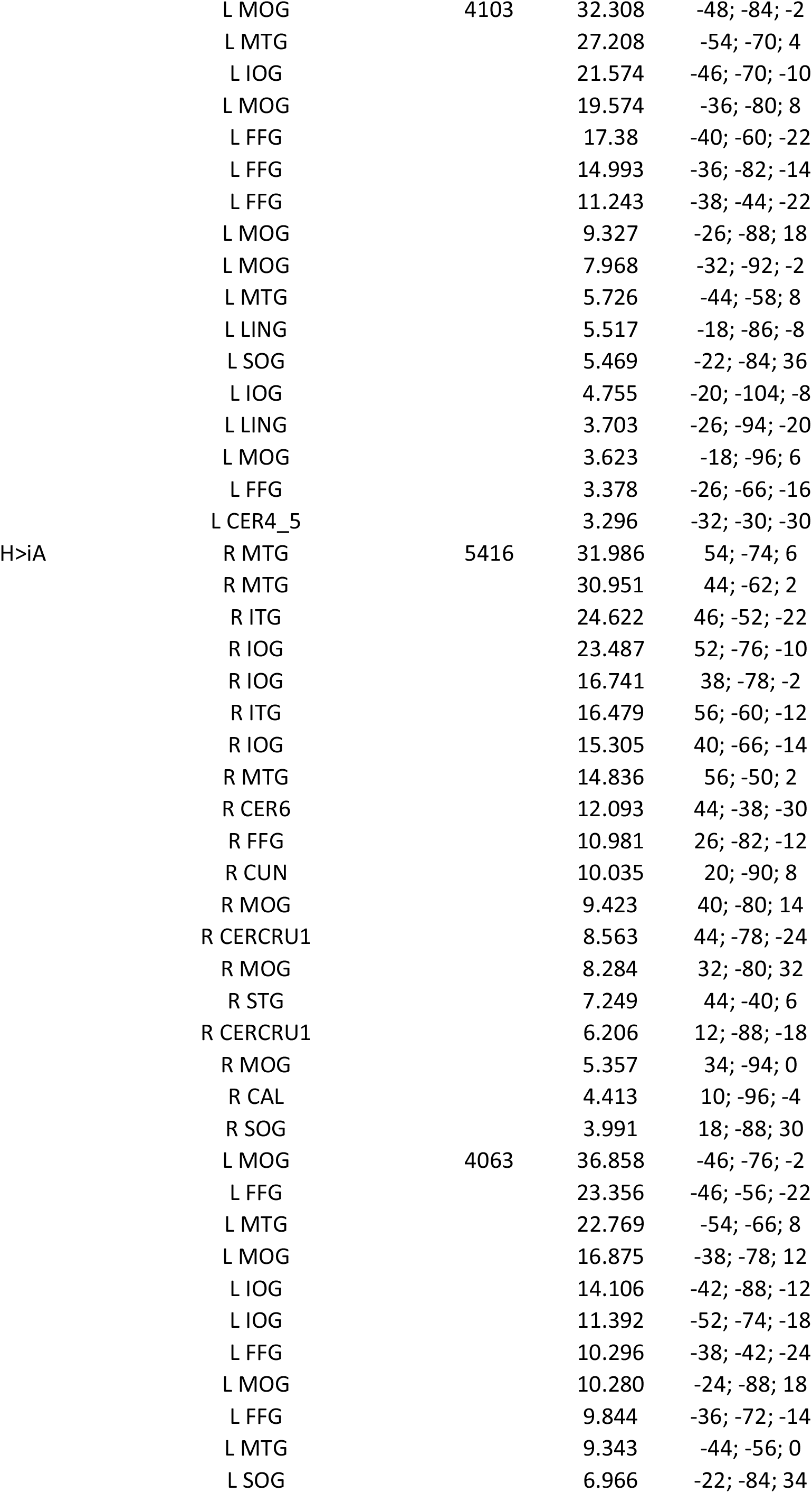
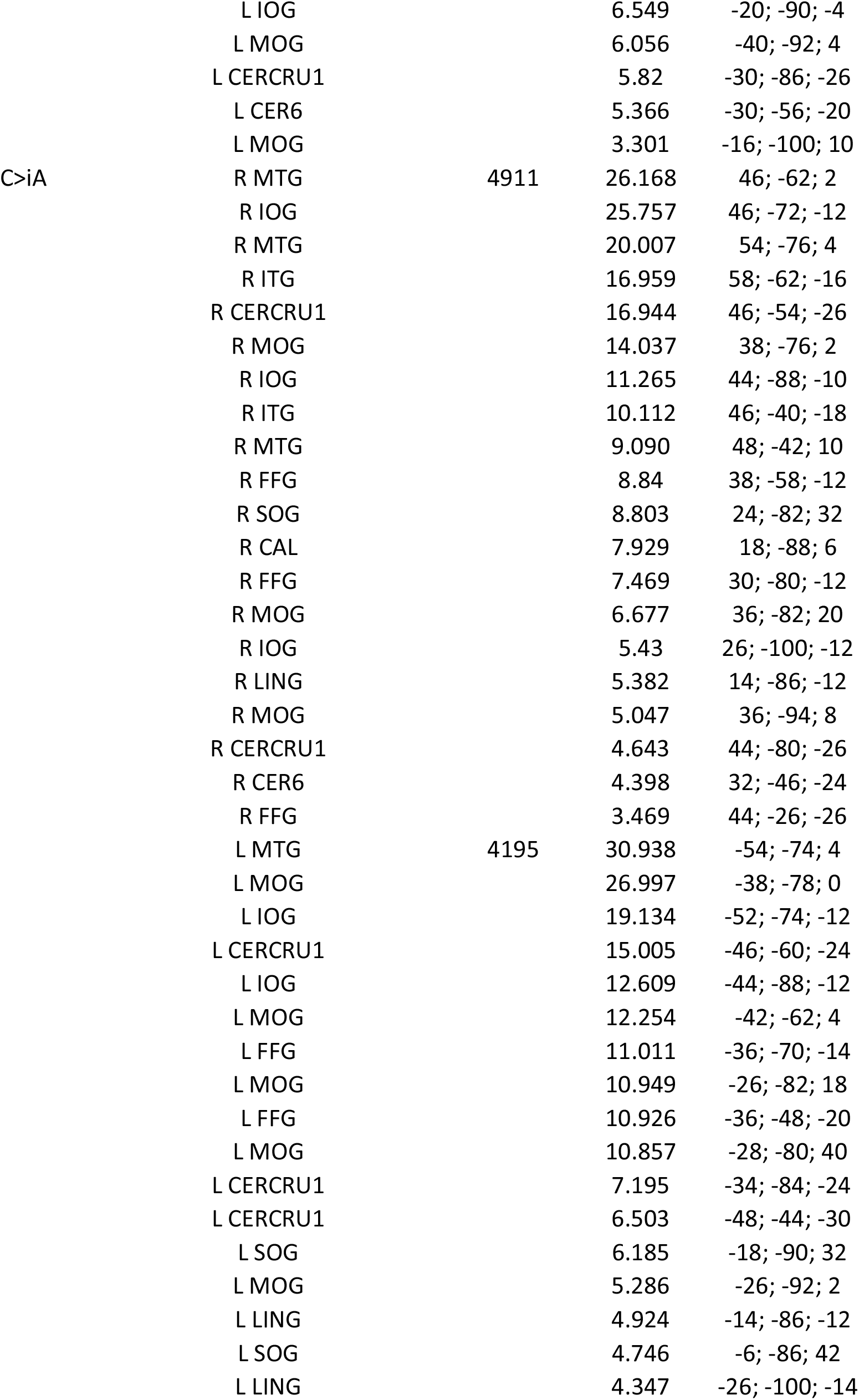
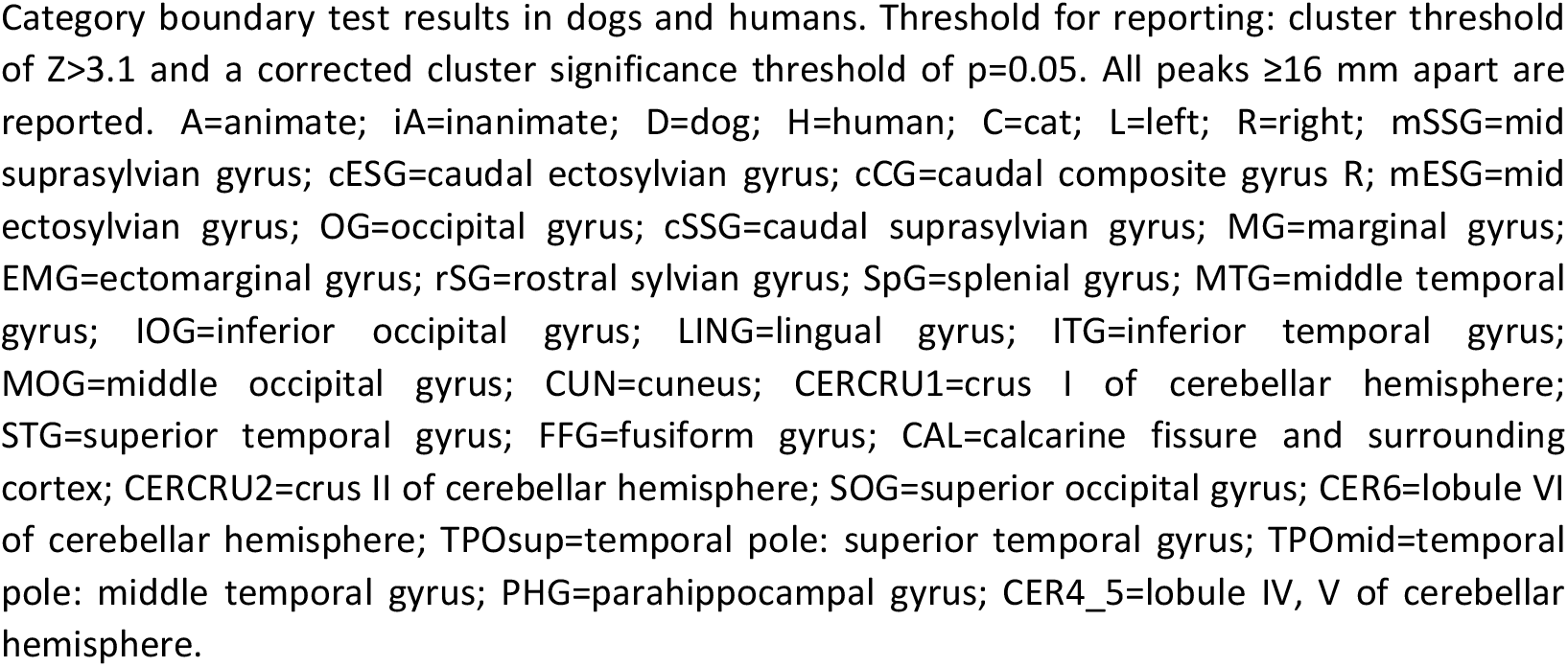

### Video legend

Supplementary Video S1. fMRI stimuli.Video shows sample stimuli representing each of four conditions: dog, human, cat, car.

Video available at Open Science Framework: https://osf.io/fzt8c/?view_only=74a9c24f13dd45078e59ad2907ec807e

